# Evaluating disease similarity using latent Dirichlet allocation

**DOI:** 10.1101/030593

**Authors:** James Frick, Rajarshi Guha, Tyler Peryea, Noel T. Southall

**Affiliations:** National Center for Advancing Translational Sciences, 9800 Medical Center Drive, Rockville, MD 20850

**Keywords:** Latent Dirichlet Allocation, Disease Similarity, Topic Modeling, Ontology, Text Mining, Online Mendelian Inheritance in Man

## Abstract

Measures of similarity between diseases have been used for applications from discovering drug-target interactions to identifying disease-gene relationships. It is challenging to quantitatively compare diseases because much of what we know about them is captured in free text descriptions. Here we present an application of Latent Dirichlet Allocation as a way to measure similarity between diseases using textual descriptions. We learn latent topic representations of text from Online Mendelian Inheritance in Man records and use them to compute similarity. We assess the performance of this approach by comparing our results to manually curated relationships from the Disease Ontology. Despite being unsupervised, our model recovers a record’s curated Disease Ontology relations with a mean Receiver Operating Characteristic Area Under the Curve of 0.80. With low dimensional models, topics tend to represent higher level information about affected organ systems, while higher dimensional models capture more granular genetic and phenotypic information. We examine topic representations of diseases for mapping concepts between ontologies and for tagging existing text with concepts. We conclude topic modeling on disease text leads to a robust approach to computing similarity that does not depend on keywords or ontology.

## 1.1 Introduction

Measures of disease similarity have been used in drug repositioning [1], drug target selection [2], and understanding disease etiology [3]. These measurements seek to make maximal use of existing biomedical research by forming hypotheses based on the similarity between a well-studied or treatable disease and one that is less well-understood. One major limitation in turning this research into insight is that the majority of disease knowledge exists in the form of unstructured free text. Since it is not easy to parse complex meaning from technical natural language, many ontologies have been constructed to classify and organize diseases.

Some researchers have used these ontologies to measure semantic or functional disease similarity [4,5] using resources like Gene Ontology [6], HumanNet [7], and Disease Ontology [8]. These approaches are based on overlap between gene sets [4] or distance in within the ontology’s hierarchy [5]. However when using an ontology to calculate similarity directly, there are a number of limitations. First, you are confined to the scope of the ontology. Most ontologies are structured in such a way that they only capture one aspect of a disease. For example, one ontology might organize diseases by affected organ system, another by disease basis, and a third by genes associated.

This means that an ontology-based similarity metric is measuring distance only in the dimension of disease captured by that ontology. In order to evaluate similarity along more than one dimension (to incorporate genetic and phenotypic information in the same measure, for example), there must exist an exact mapping between the ontologies that capture each of those dimensions. But creating such a mapping between two ontologies is often an extremely difficult task. Even expert-curated efforts at medical ontology mapping are prone to inconsistencies, false synonymy, and redundancy [9].

Some ontologies attempt to capture more than one aspect of disease, but doing so is very difficult and leads to unexpected results. For example, DO categorizes diseases by infectious agent, by organ system affected, and by mode of inheritance. Each one of these pieces of information is important, but not every category is captured for each disease. Only 71 of 8077 diseases are categorized as belonging to more than one of these categories. For example if a disease has “Genetic Disease” as an ancestor, then it is very unlikely that it also has “Disease of Anatomical Entity” as an ancestor as well, even if both might be appropriate. This does not mean that DO has incorrectly categorized any of its entries, but it means notions like node depth and ancestry have very different meanings in different parts of the ontology. This in turn makes those pieces of information less meaningful in measures of disease similarity.

Beyond purely ontology-based methods, there have also been attempts to leverage free text in measuring similarity. However these text mining approaches have still relied heavily on ontologies and controlled vocabularies, which reduce meaning and ignore context. Van Driel et al [10] calculated disease similarity using frequencies of Medical Subject Headings (MeSH) terms [11] as a way to represent disease-related text. Using this feature space, the authors represent phenotype records in Online Mendelian Inheritance in Man (OMIM) [12] as MeSH term frequency vectors. With these vectors, they compute a phenotype-phenotype similarity matrix, but report no apparent clusters within the data.

More recently, Hoehndorf et al [13] mined text from Medline abstracts to associate phenotypes with diseases using the Human Phenotype Ontology (HPO) [14] and DO. The authors use co-occurrence of DO terms with HPO terms in an abstract or title to form these associations. The paper performs a thorough analysis, using the associations to measure disease-disease similarity and to predict gene-disease associations based on phenotypic similarity. The approach demonstrates the ability to cluster diseases based on this similarity metric. However the method is fully reliant on matching curated terms or keywords from the ontologies involved to the abstracts and it limits itself to studying phenotypic similarity.

Our method seeks to increase the flexibility of this analysis by using all of the text associated with a disease instead of simply counting keywords. We use Latent Dirichlet Allocation (LDA) [15], an unsupervised probabilistic method of learning topics associated with text. In the biomedical space, LDA has been used to analyze electronic health records [16, 17], drug labels [1], and adverse event data [18].

One benefit of using LDA is removing the strict reliance on a given ontology. Instead of learning only from the keywords which map directly to an ontology class, LDA can use a vocabulary more tailored to the corpus on which it is trained. Additionally, LDA can form associations from multiple types of information at once. Topics include a mixture of genetic or phenotypic information as well as any other clinically relevant characteristics included in the text. It is especially useful that the output of the model is interpretable and can be easily inspected.

We learn numeric vector representations of diseases as mixtures of topics from free text. We use these representations to calculate disease similarity, but we also explore their usefulness in concept normalization and mapping concepts between ontologies. We evaluate our similarity metric using DO as ground truth and we show that the representations can be used for tagging biomedical corpora.

## 1.2 Methods

The goal of this process was to quantify similarity between diseases by leveraging free text disease descriptions. The approach first creates a topic model using LDA on textual entries in a disease database, then uses an orthogonal ontology as ground truth to validate the model (Figure 1).

**Figure 1.**
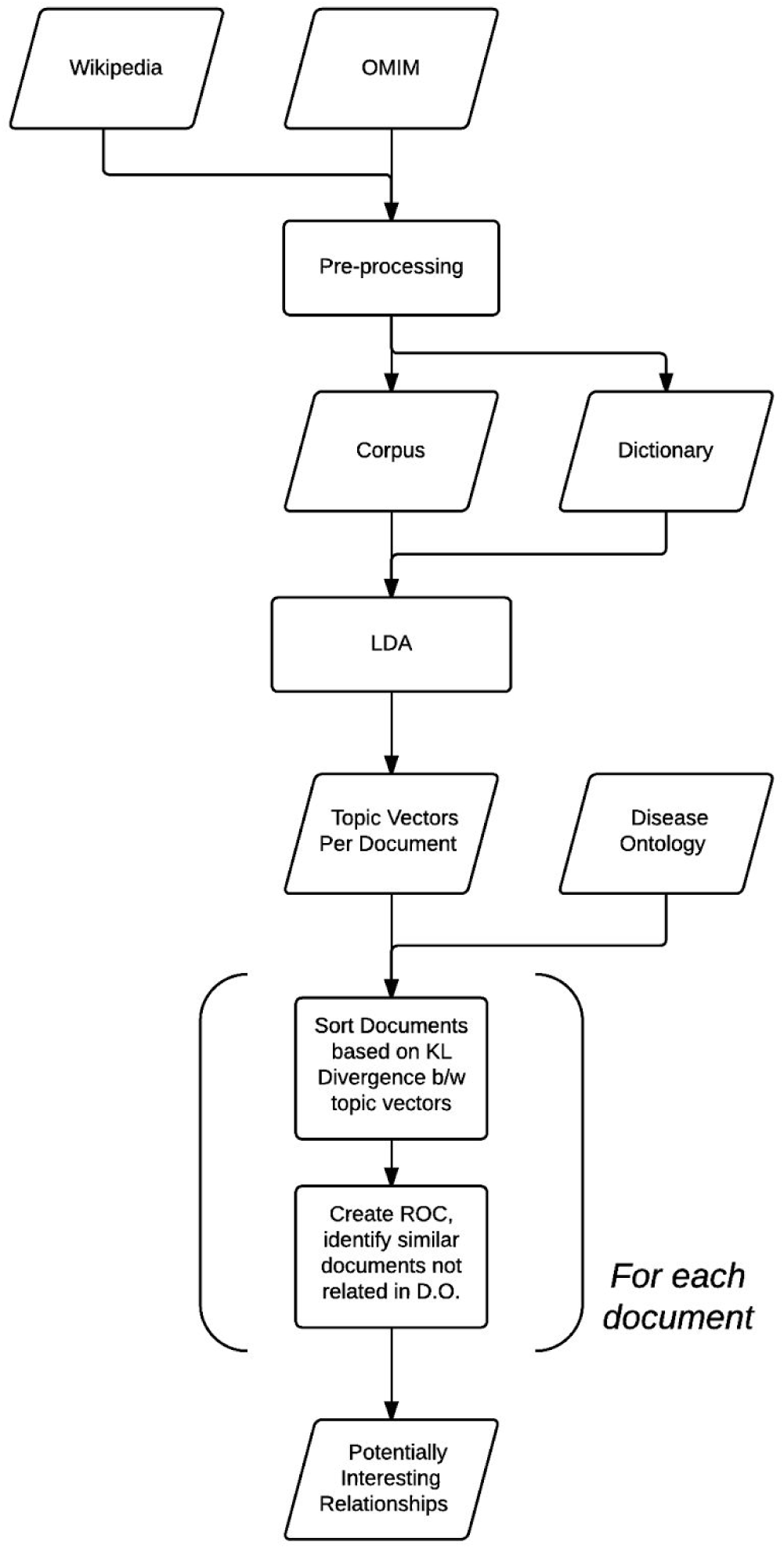
Overview of the proposed methodology and validation

### 1.2.1 Latent Dirichlet Allocation

LDA is a generative bag-of-words approach to topic modeling. It relies on the assumptions that the order of words within a document does not matter and that a collection of documents were generated by sampling from an underlying distribution of topics. The algorithm outputs a distribution over words in its dictionary for each topic. As the algorithm iterates over the documents, words are clustered together into topics and documents are assigned the topics most likely to have generated them. The LDA implementation employed in this paper is an online version of variational Bayes LDA [19].

The theoretical basis for LDA is fully described in [15]. Briefly, the algorithm works by iteratively performing the following steps:

□ Assign each word *w* in each document *d* to a random topic

  □ For each document *d:*
  
    □ For each word *w*:
    
      □ For each topic *t*:
      
        □ Calculate probability of topic *t* given document *d* using current word-topic assignments within *d*
        □ Calculate probability of word *w* given topic *t* using the current contribution to topic *t* from word *w*.
      □ Reassign word *w* to topic *t* with the probability distribution *p*(*t/d*)*\*p*(*w/t*) for all topics
  □ Repeat

The computational challenge is the calculation of the posteriors over word and topic probabilities. The algorithm first makes the simplifying assumption of dropping the dependencies that make this inference intractable. With this simplification it becomes possible to find an approximation of each posterior, q(z, *θ*) from a family of possible distributions with new free variational parameters γ and ϕ. The algorithm finds values of these free parameters which minimize the Kullback Leibler divergence [20] between the approximation q(*θ*, z | *γ, ϕ*) and the true posterior, p(*θ*, z |w, *α*, *β*).

### 1.2.2 Document collection

To demonstrate the method we used as our corpus the 4558 descriptive records in OMIM as of September 2015. The training corpus for the model also included the text from the 1339 Wikipedia articles that directly called out an OMIM identifier in our document set. This was done to increase the amount of text about each disease the model sees and to improve applicability of the model to non-OMIM text. The Wikipedia documents were used to improve training but were not evaluated in our validation step. In all cases we only employed English language documents.

### 1.2.3 Dictionary generation and document pre-processing

After collecting the relevant documents, we performed a series of pre-processing steps to prepare the documents for use in LDA. First the documents were stripped of references, headers, and other source-specific text in order to minimize bias across sources. Documents were split on whitespace/punctuation and lowercased to form individual word tokens. Stopwords were removed from the documents using a dictionary provided in the Natural Language ToolKit (NLTK) [21]. English words were stemmed using NLTK’s implementation of the Porter Stemmer. Other words and tokens were left unprocessed to avoid stemming scientific terms unnecessarily.

The top *V* most frequently occurring words in the corpus comprised the dictionary used by the LDA algorithm. Dictionary size, V, was a parameter varied in optimizing the model. After pre-processing, documents are essentially just a list of pointers to entries in a dictionary. Anything outside the dictionary is ignored, and word order does not matter during construction of an LDA model.

### 1.2.4 Defining disease similarity

By associating text with each disease we want to consider, we can now assign topic vectors to each disease. In other words, a disease is defined in terms of the contribution of each of the topics computed by the LDA model. LDA is a generative method which assumes that documents are generated by first picking a topic *t* from a distribution over all possible topics, then choosing a word from a distribution over all words given that topic. The process of training the LDA model is determining the distributions most likely to have given rise to the observed documents. After training, the topic distribution for a given document represents the parameters of the Dirichlet distribution over topics which generated that document. The output of this procedure is a real-valued vector of length *K* (where *K* is the number of topics) for each document which represents the theoretical contribution of each topic to the document’s generation.

By comparing the distance between the topic vectors learned from two documents, we have a measure of how similar we believe those documents’ diseases to be. Concretely, we assume that the model is approximating topics that represent actual concepts in the disease space. Thus if two documents appear to have been generated from the same set of topics, their diseases may demonstrate shared etiology. We use the KL divergence as our distance metric, as it is used as a measure of similarity between two distributions (in this case a Dirichlet distribution over topics). More specifically, it measures how much information is lost if you use the topic distribution of the result document to approximate the distribution of the query document.

## 1.3 Results

### 1.3.1 Disease Ontology Evaluation

When it comes to evaluating the performance of our similarity measurements, there is no obvious gold standard because disease similarity lacks measurable ground truth. Hoehndorf et al. [13] evaluate their model by determining to what extent their measure of similarity between human disease phenotype and mouse phenotype predicts mouse disease genes. But that metric does not translate to our methodology due to a lack of free text descriptions of mouse models. Cheng [4] and Mathur [5] both use a very small, curated set of disease relationships as their test set, but of the roughly 70 relations these two works use, only 5 can be linked to one of our OMIM records. Our solution was to use relations within Disease Ontology as a ground truth measure of similarity.

We considered two DO entries to be related if they shared a common ancestor within 3 generations. Of the 1613 records that had a DO mapping, 302 of them were only children of “Autosomal Recessive Disease” or “Autosomal Dominant Disease.” The extremely shallow nature of this categorization led to a number of ground truth relations between otherwise dissimilar diseases. For the analysis below, we excluded these records (leaving 1311 records), but in general including them had little effect on the process when our topic number was low, but negatively impacted performance in higher dimensions. This makes intuitive sense, as higher dimensionality will lead to more granular topics, which will do a worse job at capturing similarity if a larger percentage of ground truth relations are high level.

For each of the 1311 OMIM documents (our “query”), we sorted all other OMIM records in the set based on the KL divergence between the query document’s topic distribution and every other document’s topic distribution. By calling the first *n* most similar documents in our set “positives” and all the other documents in the set “negatives” we scored each document according to whether or not the “positive” documents were in fact related to the query document according to DO. By varying *n* (the number of positive documents returned), we built a Receiver Operating Characteristic (ROC) curve for each document and computed the Area Under the Curve (AUC) (Figure 2).

**Figure 2.**
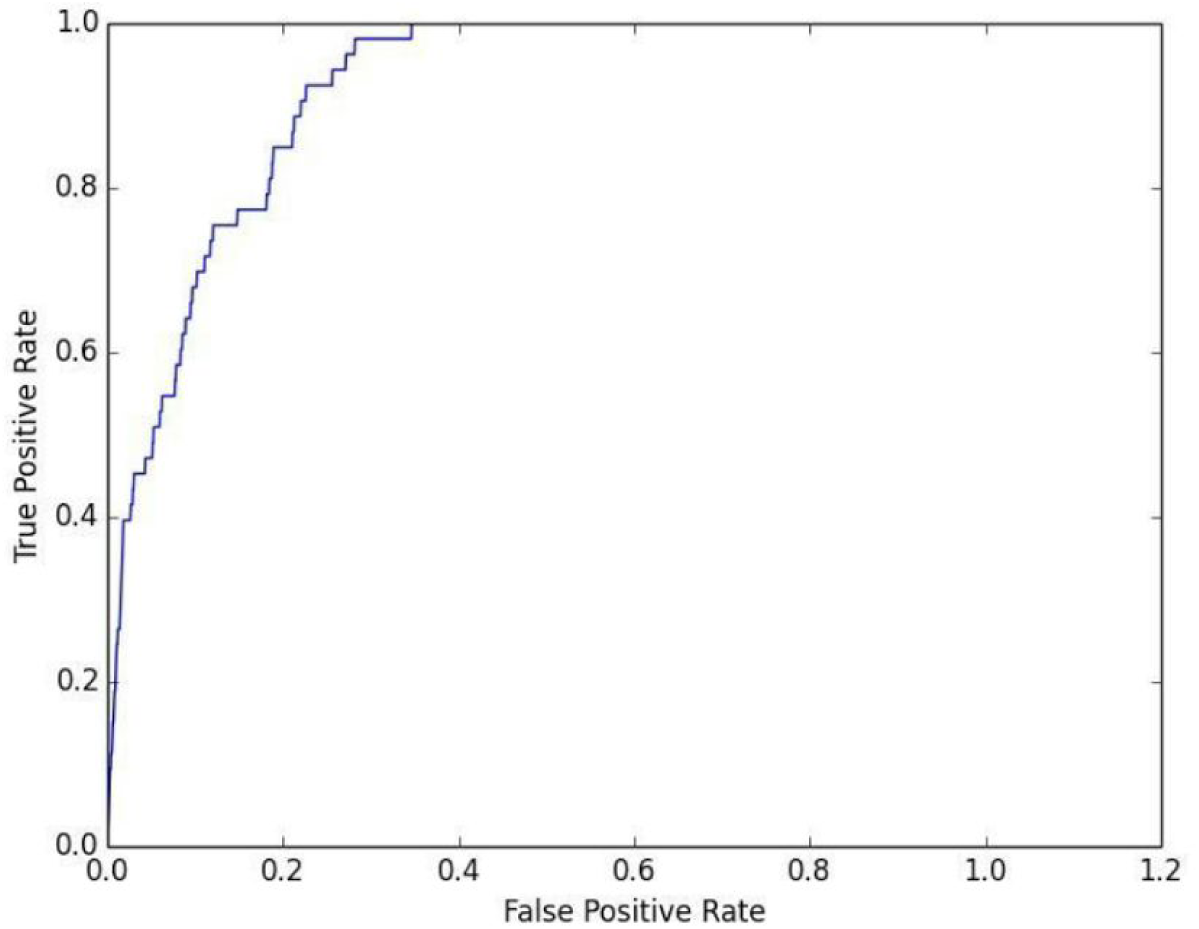
ROC Curve for OMIM_#230900 Gaucher Disease Type II on the Disease Ontology categorization task

We optimized our model’s parameters (Figures 3, 4) and hyperparameters (Figure 5) using the mean AUC and the count of AUCs above 0.90. The mean AUC of a model with 25 topics and a dictionary of 7,000 words was 0.80 ± 0.01. The mean number of documents for which the model exhibited an AUC above 0.90 was 353 ± 63 (out of 1311). The model performs best with a topic number between 20 and 50 and a dictionary size of 5000 to 7000 words. Higher dimensionality slightly impairs performance on this task while increasing the granularity of the topics learned.

**Figure 3.**
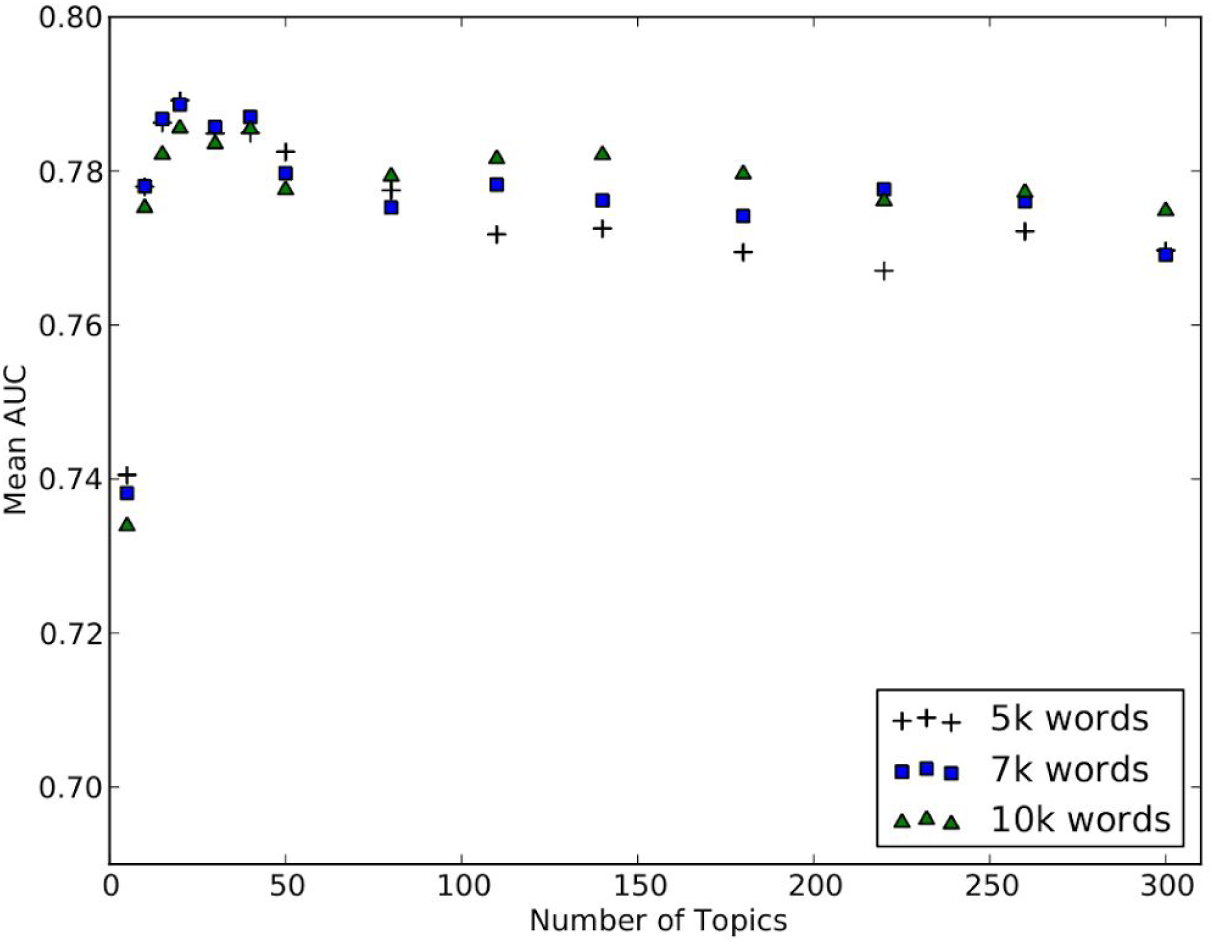
Mean AUC of LDA model on Disease Ontology task for various topic number and dictionary size combinations

**Figure 4.**
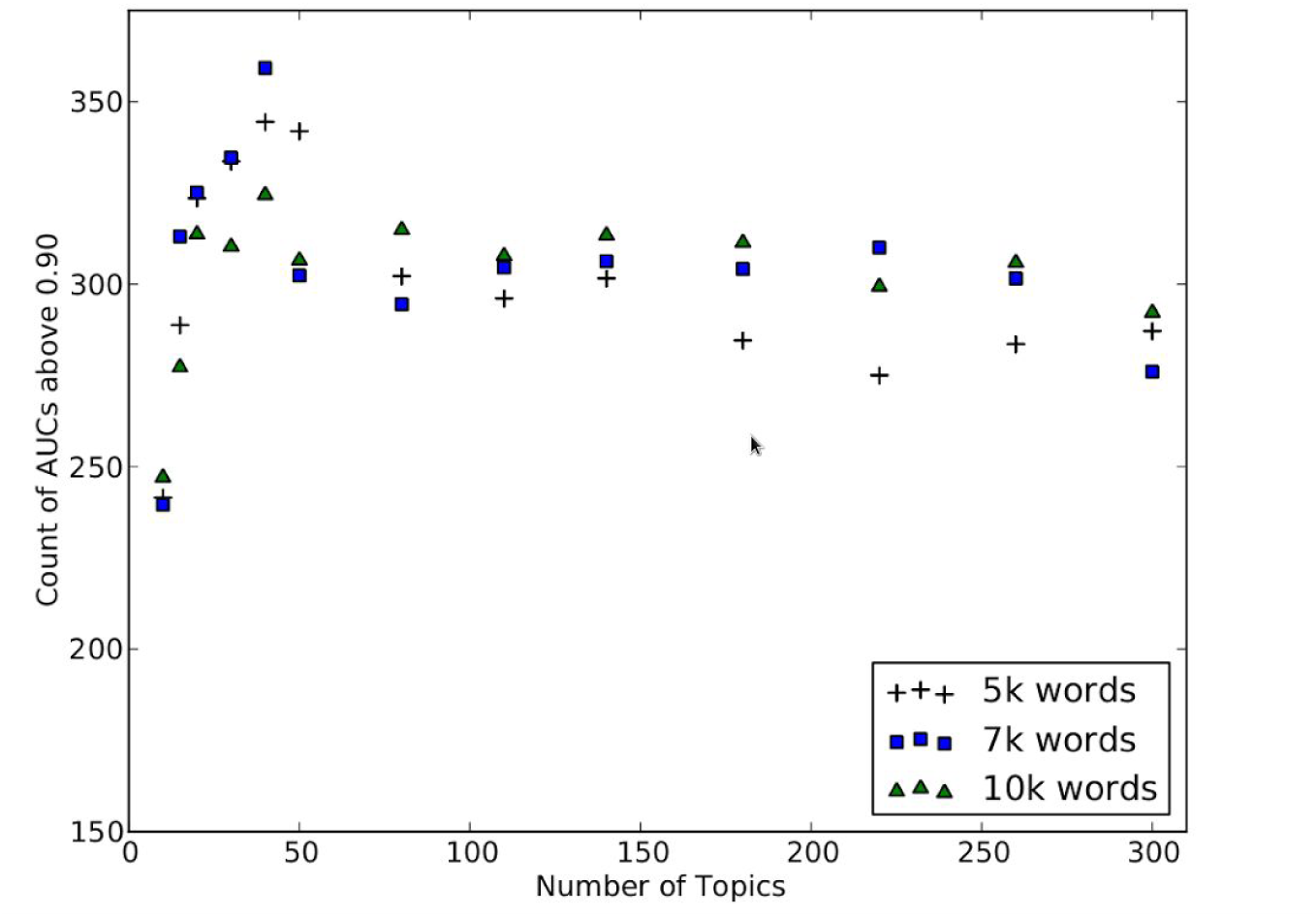
Count of records with AUC > 0.90 for LDA model on Disease Ontology task for various topic number and dictionary size combinations

**Figure 5.**
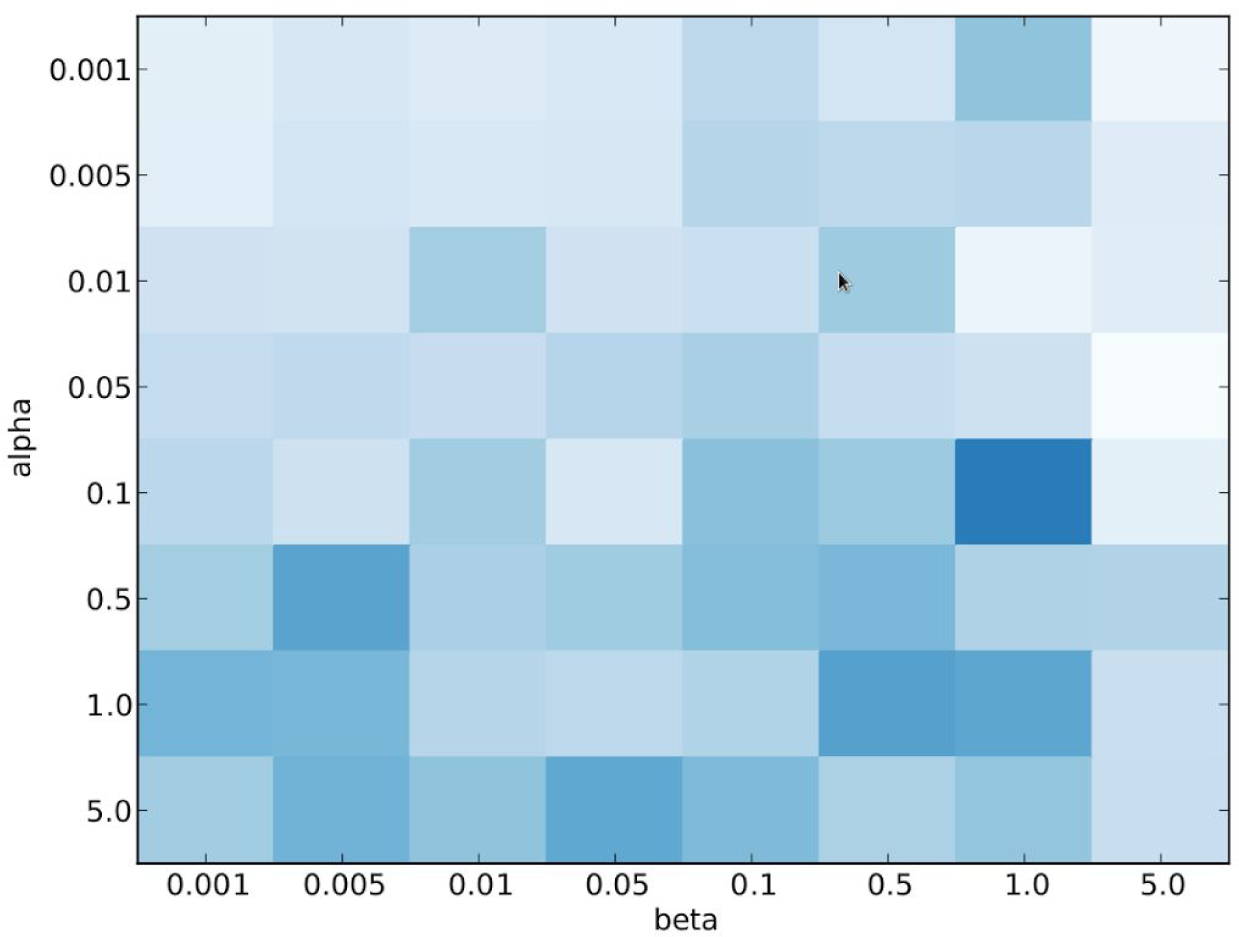
Heatmap of mean performance of LDA model on Disease Ontology task in hyperparameter space

The most similar documents to most queries shared obvious relations with the query document (Table 2, Figure 6), though these are not always the kinds of relations captured in DO. Interestingly, the model often learns that different variants of the same disease class are not the most similar diseases. For example OMIM230800 Gaucher Type I is less related to OMIM230900 Gaucher Type II than it is to OMIM607625 Niemann Pick Type C2.

**Figure 6.**
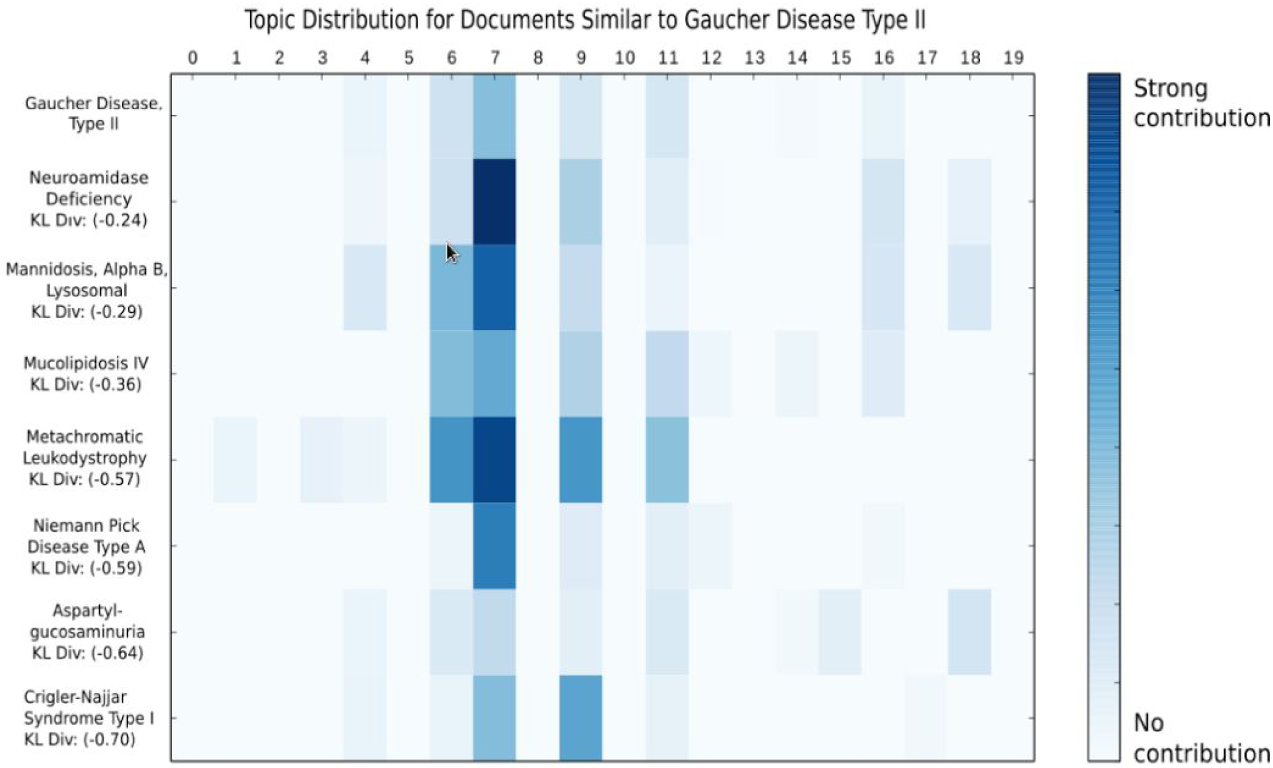
Example topic distribution for documents most similar to Gaucher Disease Type II determined by an LDA model with 7000 words and 20 topics

This may be a weakness or strength of the model depending on the application. It shows that the similarity is based on a latent representation of the diseases rather than on the presence of a few keywords. In this example, we can see that the primary topic to which these records belong contains many relevant words. Obviously it contains “lysosomal” and “storage” as these are all lysosomal storage diseases. But very strongly associated are the words “lipid(s),” “level(s),” “accumulates/accumulation,” “acidic,” “lipoprotein,” and “deficient/deficiency.” The topic also contains obvious keywords such as “Gaucher” and “Neimann” but can represent a lysosomal storage disorder in a much richer fashion than simply keyword matching.

A useful result of this methodology is that we learn topic vectors for many OMIM diseases that do not have a DO mapping. In these instances, our similarity metric served as a way to place an unmapped OMIM disease within the DO. For example according to our methodology, the document most similar to OMIM126600, Doyne Honeycomb Degeneration of Retina (which does not map to an existing DO record), is Stargardt Disease which DO categorizes as an “Age-Related Macular Degeneration.” The other top documents were Basal Retinal Drusen and Vitelliform Macular Dystrophy, both characterized by DO as one level higher, a “Degeneration of Macula” (Figure 7). Examination of the nearest neighbors of a given document seems to be a functional way to suggest node location within a target ontology.

**Figure 7.**
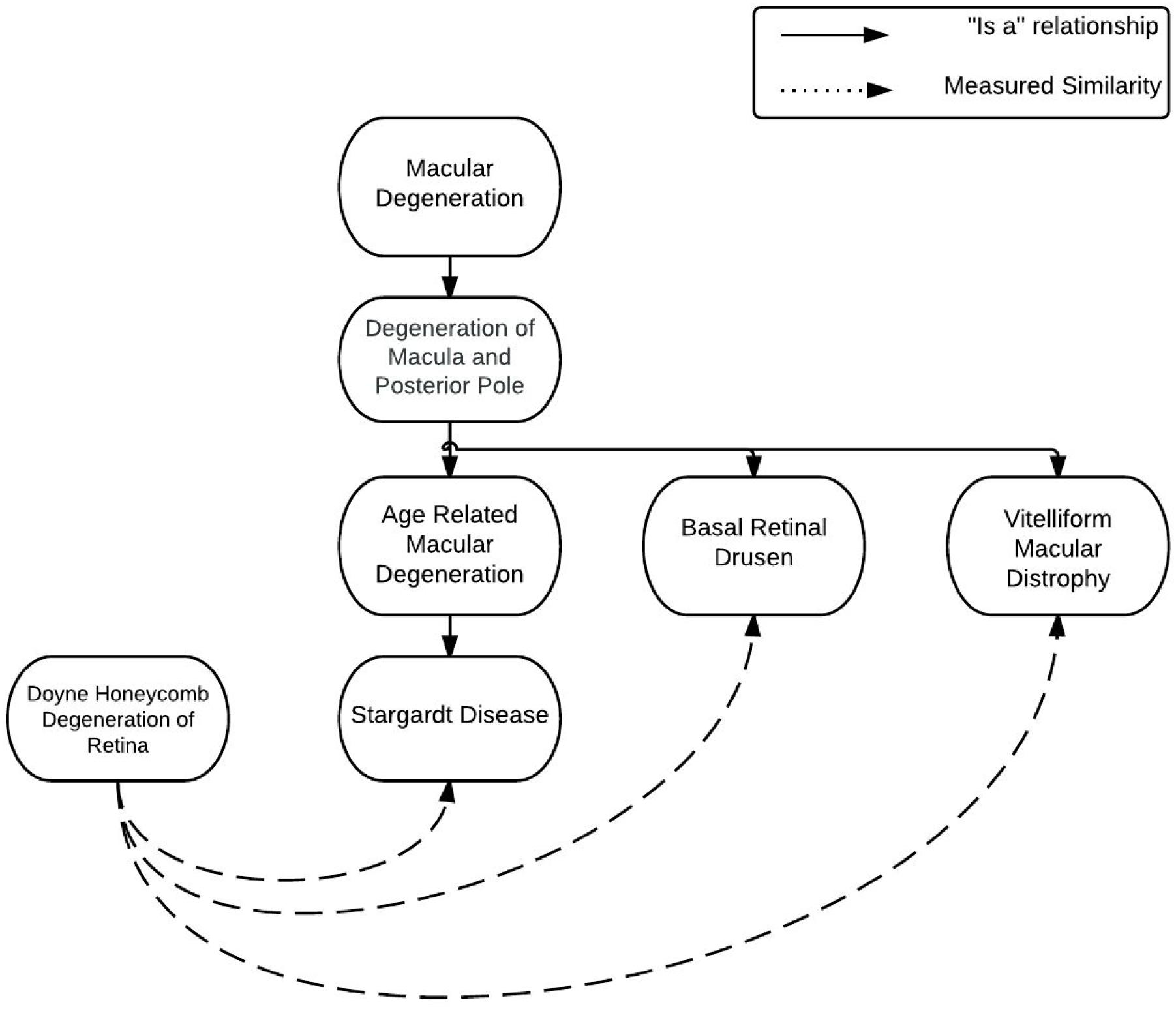
Demonstration of the utility of the LDA method for placing unmapped concepts within an existing ontology

It is important to note that the DO evaluation metric is orthogonal to the training of the model. LDA is unsupervised, so in effect we “hold-out” all of the ground truth relations each time we train a new model, then evaluate on all those held-out relations. The purpose of the AUC measurement is not to determine how well the model would generalize to new examples, but rather to validate our measure of similarity.

### 1.3.2 NCBI Disease Corpus

The NCBI Disease Corpus [24] is a manually curated corpus of 793 titles and abstracts tagged with the disease concepts mentioned therein. The tags reference a MeSH or OMIM concept. The dataset is designed to be a gold standard in disease concept normalization and is split into training and test datasets. Traditional approaches generally rely on Named Entity Recognition (NER) to extract terms before normalization [25]. Using a state-of-the-art NER application allows a fairly simple concept normalization scheme to achieve high precision.

We tested LDA disease vectors for the combined task of NER and concept normalization in the NCBI Disease Corpus. We first constructed a document for each disease concept in the training set. Each document consisted of the text from the abstracts in which it was mentioned in the training set. We then performed LDA on these documents to generate a topic vector for each concept. Then we showed the model each of the abstracts in the test set to learn their topic vectors. For each test set topic vector, we compared with every topic vector from the training set and ordered them by topic similarity. This approach achieved a mean AUC of 0.754 on the task (Figure 8), despite the fact that only 144 of the 200 classes in the test set were mentioned in the training set. When a label in the test set does not appear in our training set, we have no topic vector for it, and therefore we will not do better than random guessing.

**Figure 8.**
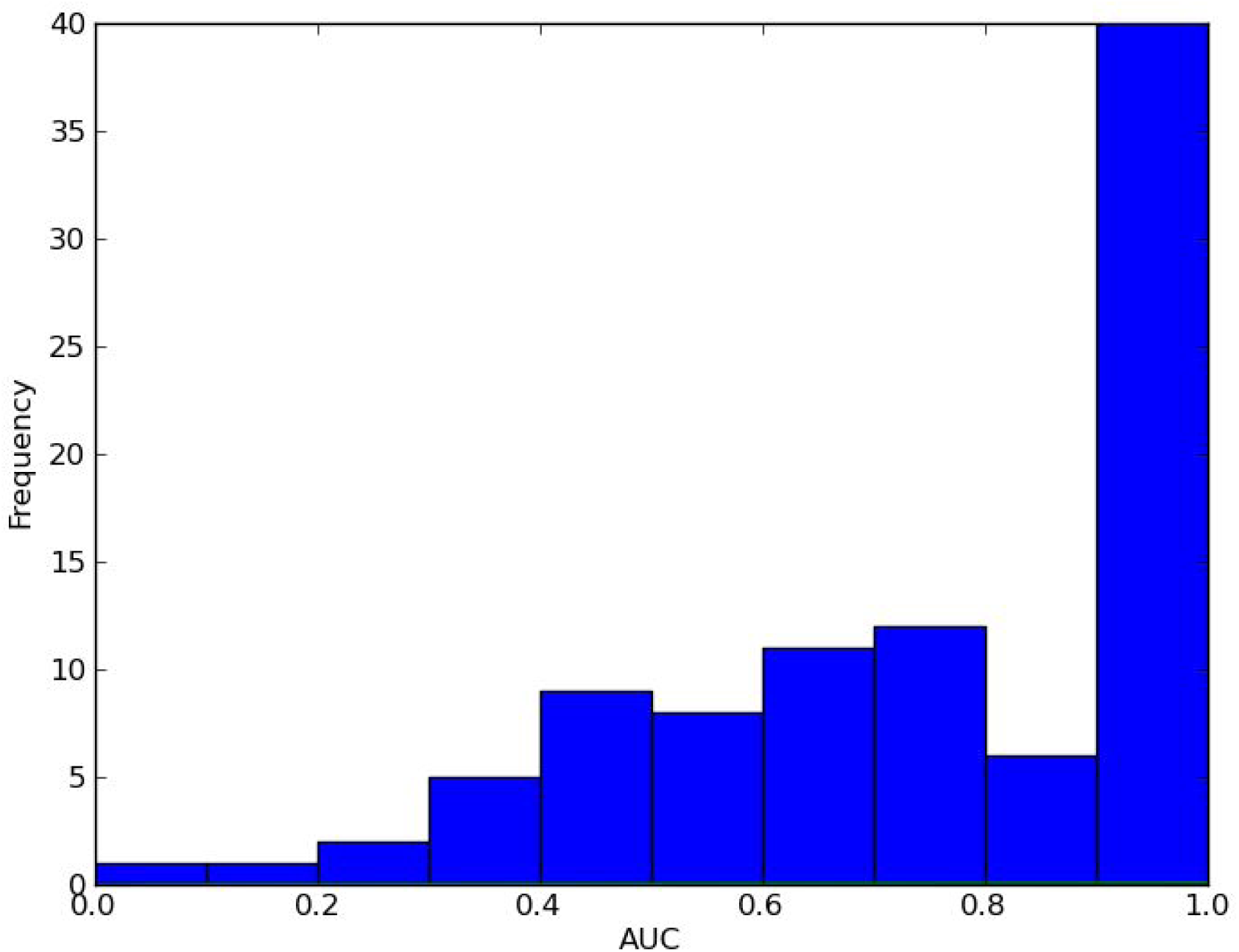
Distribution of AUCs for LDA model on concept identification and concept normalization task using NCBI Disease Corpus

### 1.3.2 Parameter Selection

The method is dependent on a number of tunable parameters (topic number, dictionary size, Dirichlet hyperparameters). As a result, it is possible to generate many different models from the same data. To tune the possible parameters of the model, we evaluated the performance of the method on the DO task for a wide range of values (Table 1). We evaluated both mean AUC across the document set as well as count of AUCs above 0.90.

**Table 1.**
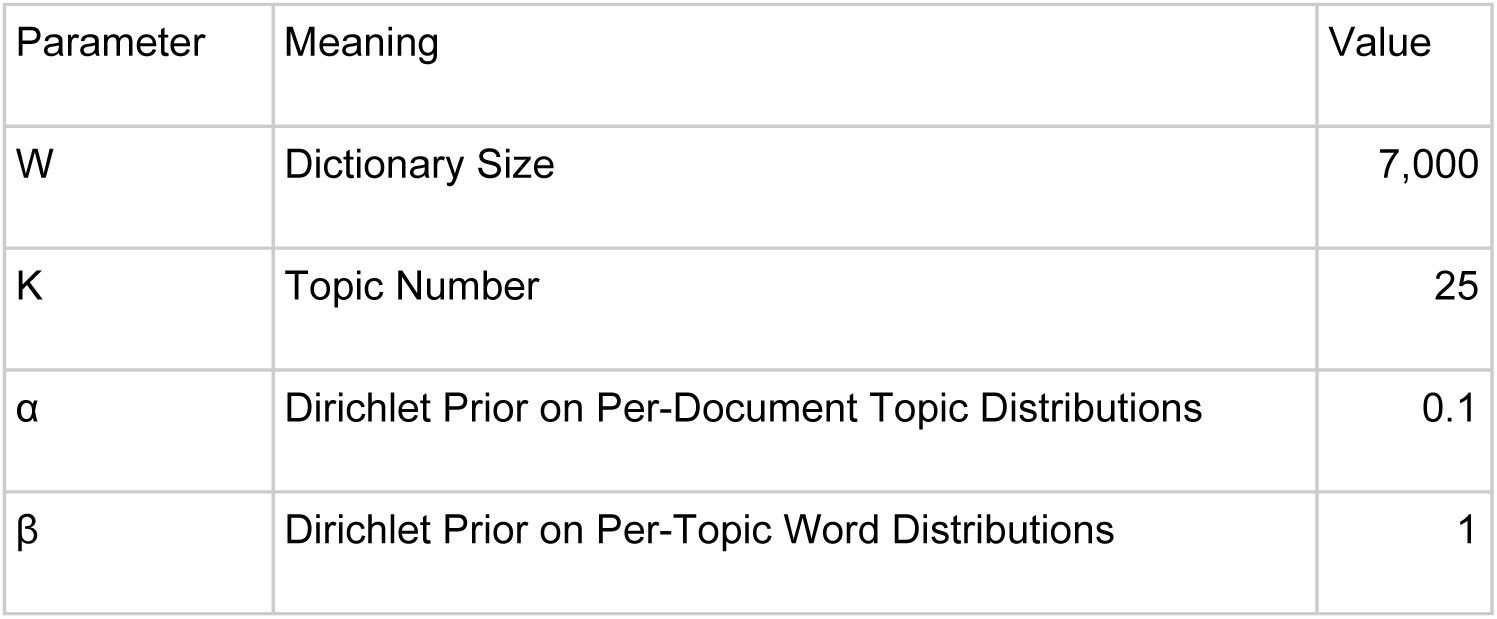
Values parameterizing the LDA model that performed optimally on the Disease Ontology categorization task

| Parameter | Meaning | Value |
| --- | --- | --- |
| W | Dictionary Size | 7,000 |
| K | Topic Number | 25 |
| $\alpha$ | Dirichlet Prior on Per-Document Topic Distributions | 0.1 |
| $\beta$ | Dirichlet Prior on Per-Topic Word Distributions | 1 |

**Table 2.** Sample of query documents with AUC > 0.90 and their 5 most similar records

| Query Document | Most Similar Documents |  |  |  |  |
| --- | --- | --- | --- | --- | --- |
| Machado-Joseph Disease | Spinocerebellar Ataxia 1 | Frontotemporal Dementia | Spinocerebellar Ataxia 2 | Spinocerebellar Ataxia 6 | Dentatorubral-Pallidoluysian Atrophy |
| Macular Dystrophy, Vitelliform, 2 | Oguchi Disease 2 | Leber Congenital Amaurosis 6 | Doyne Honeycomb Retinal Dystrophy | Night Blindness, Stationary, Type 1F | Retinitis Pigmentosa |
| Gaucher Disease, Type II | Gaucher Disease, Type III | Gaucher Disease, Saposin C Deficiency | Niemann-Pick Disease, Type C2 | Gm1-Gangliosidosis, Type III | Gm1-Gangliosidosis, Type II |
| Night Blindness, Congenital Stationary, Type 1B | Oguchi Disease 2 | Leber Congenital Amaurosis 6 | Retinitis Pigmentosa 42 | Cone-Rod Dystrophy, X-Linked, 1 | Night Blindness, Stationary, Type 1F |
| Osteogenesis Imperfecta, Type III | Osteogenesis Imperfecta, Type IV | Osteogenesis Imperfecta, Type I | Achondrogenesis, Type IB | Ehlers-Danlos Syndrome, Type IV | Osteopetrosis, Autosomal Dominant 2 |
| Tay-Sachs Disease | Sandhoff Disease | Niemann-Pick Disease, Type C1 | Gaucher Disease, Type II | Gaucher Disease, Type III | Niemann-Pick Disease, Type B |

### 1.3.3 Control Experiments

Given the complexity of the LDA methodology, we investigated simpler alternatives. For comparison, we applied a similar approach applying Principal Component Analysis (PCA) to normalized term frequency (TF) vectors for each document in the corpus. We generated a *D* × *V* matrix of word frequencies, normalized both within document and within features. After performing PCA, we kept as many principal components as we had topics. We then constructed ROC curves for the DO mapped documents, as done above for LDA, based on distance between TF PCA vectors. In lower dimensions, this control approach performed similarly to LDA, but with important differences. Since LDA is probabilistic and unsupervised, we see variance in its performance. Thus by guiding it, (either by simply adding keywords to each document or by using one of several existing supervised approaches to LDA [22,23]), we can achieve performance improvements. Secondly, as we increased dimensionality the additional principal components helped performance less than the additional topics. This is a result of the nature of principal components, each of which captures successively less variance.

Notably, the number of principal components at which our performance measure leveled off was similar to the optimal number of topics in LDA (Figure 9), perhaps related to the number of dimensions required to capture the variance in the DO hierarchy. The TF approach also leverages all the free text associated with a disease and performs almost as well as LDA in low dimensionality. However it does so at the cost of interpretability. Examining the word distribution over each topic provides an easy way to understand what is driving each latent feature.

**Figure 9.**
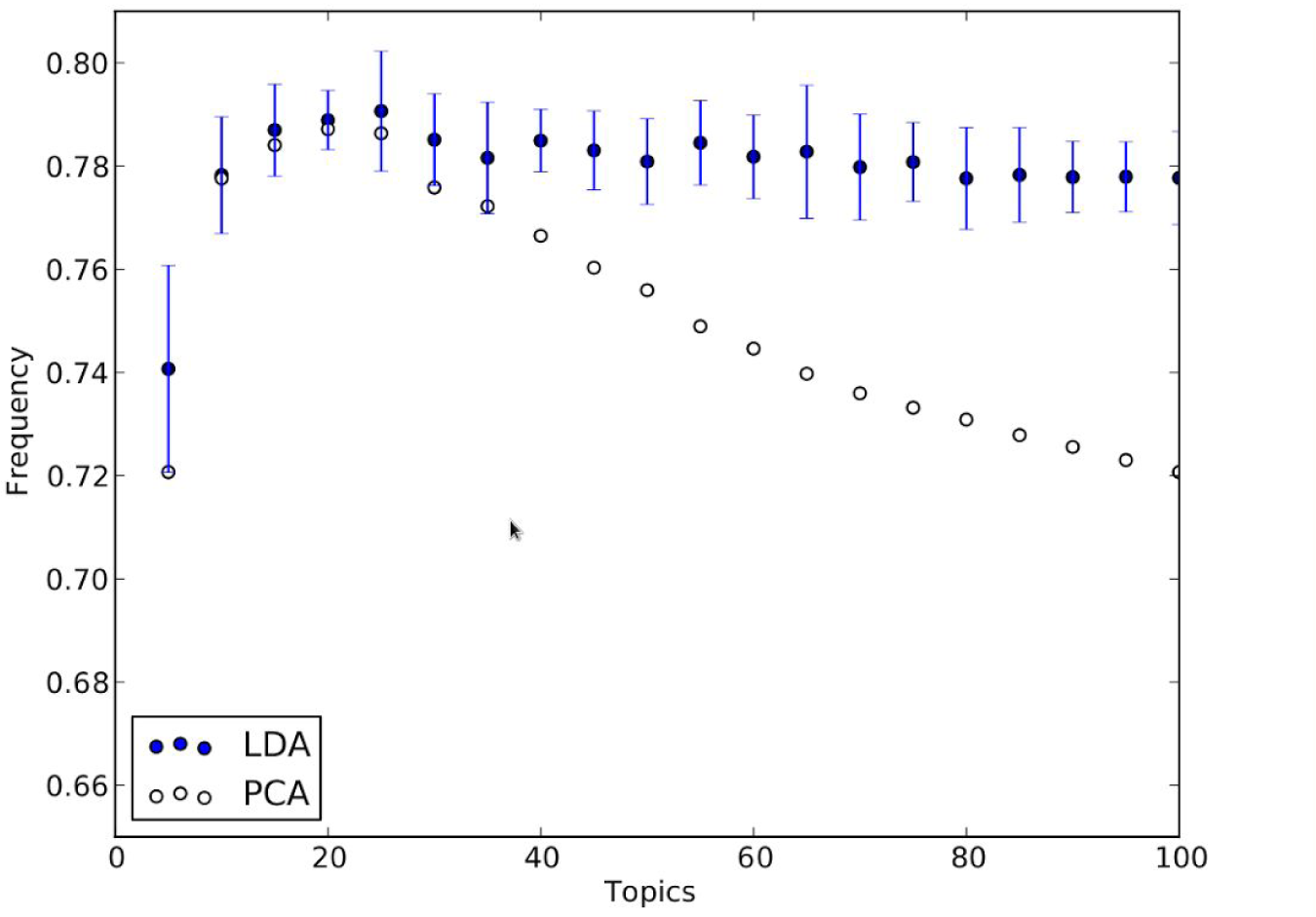
Mean AUC on DO task for an LDA model with a 7,000 word dictionary vs PCA keeping the same number of principal components as topics.

To ensure that the metric itself was not biased, we shuffled the labels of the OMIM records thus randomizing the connections between documents within the DO. We then performed the same process and as expected the resulting distribution of AUCs was tightly centered around 0.5 as expected (Figure 10).

**Figure 10.**
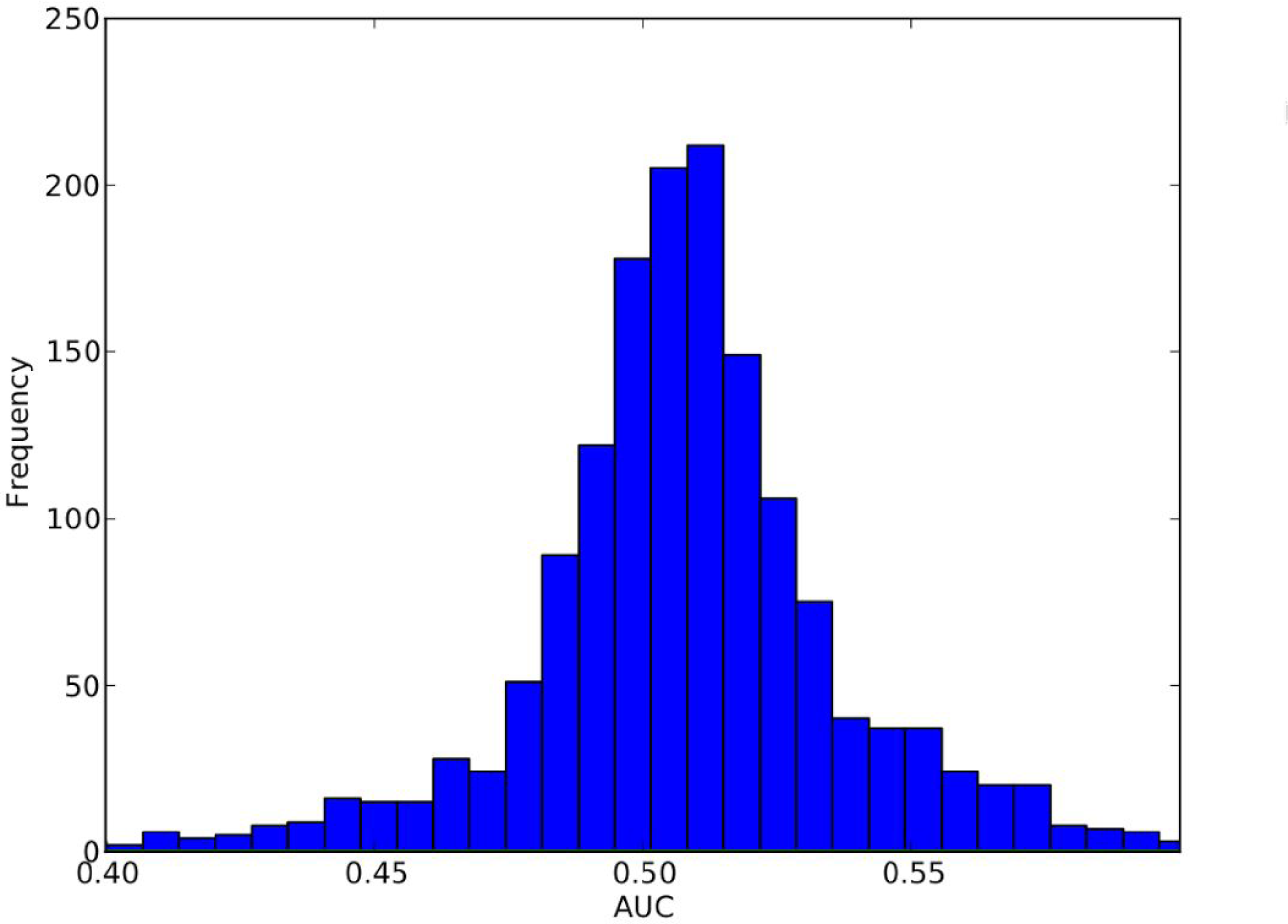
Distribution of AUCs on Disease Ontology categorization task with shuffled Disease Ontology IDs.

### 1.3.4 Interpretability

LDA offers the advantage of having an interpretable output; once the model is trained, we have a *K* × *W* matrix that represents the distribution of W words over K topics. This offers the ability to look for meaning within topics and to see what words influence the model most.

For small topic number *K*, we found that the topics tended to represent more anatomical and phenotypic information. Inspecting the topics, we could see topics describing nephrology, cardiovascular problems, neurology, blood, and tumors. At slightly larger values of *K*, we saw topics start to represent high level classes of diseases. One topic contained the terms “Huntington,” “ALS,” “Parkinsons,” and “Alzheimer.”

At much larger K, the topics converged very differently and became more specific to individual diseases. In a run with 200 topics, one topic contained the terms, “PKU,” “PAH,” “phenylalanine,” “phenylketonuria,” “PHE.” This is a particularly illustrative case as PAH is an overloaded abbreviation in the disease space; it means both “Phenylalanine Hydroxylase” and “Primary Arterial Hypertension.” However the model learns to correctly associate PAH with this topic because it considers each term in the context of the surrounding document.

The top words in each topic are provided for two different size models as well as a visualization of the topics (Additional Information 1-3).

## 1.4 Discussion

It is difficult to arrive at a precise quantitative evaluation of this methodology due to the lack of ground truth similarity scores between diseases. The DO categorization task serves as an indication that the method learns reasonable topics which correspond to a human understanding of disease. But an AUC of 1.0 on the task would simply mean we could recreate DO’s structure, not necessarily that the similarity metric was optimal. Rather, we expect that the topics generated using this approach are learning a richer representation of disease space than the structure of any one ontology can capture.

This approach is affected by the availability of relevant, tagged disease text. Here we restricted the task to evaluating descriptive OMIM records, but to increase the scope of the application we would require text that was well-tagged within a broader database. This can take the form of indexed scientific literature, Wikipedia entries, or curated disease descriptions. With a larger corpus, the model would be able to learn an even richer representation of each disease.

The availability of disease descriptive text is particularly limiting in the NER and concept normalization tasks. Using abstracts that only mention a disease (as opposed to describing it) as a seed for the topic representation of a concept makes it easy for our model to lack specificity. Another limitation of our approach to tagging is that we only evaluated text at the abstract level. If we had added a measure of sentence level similarity, we likely would have seen better results but at the expense of runtime and simplicity.

Adding a K-Nearest Neighbors methodology as a final step after generating LDA topic vectors could be a formalized way to use this approach to extend or improve an ontology. In the case where an ontology is too flat or lacks valuable, observable relationships, it may help to use document similarity to improve the ontology’s structure. For example, DO has many nodes whose only parent is “Autosomal Dominant/Recessive Disease.” While this is correct, it does not convey much information about the disease. By evaluating the parents of documents that have similar topic distributions, it might be possible to break out these large clusters into more specific ones.

The procedure described here can be extended in a number of ways. First, for modeling the hierarchy inherent to an ontology, a hierarchical implementation of LDA could be used. It would also be easy to take advantage of the structure of an ontology to add terms identifying a document’s parents to the associated text, thereby guiding the learned topics. If using a larger corpus, the method would also likely benefit from including n-grams in the dictionary.

Additionally, once a model like this has captured a latent representation of an entity from an ontology, the topic distribution can be used to identify similar text from any other source. This presents a robust alternative to keyword matching for tagging scientific literature.

## 1.5 Conclusions

The goal of the approach outlined above was to learn vector representations of diseases from their textual descriptions, then use those representations to measure disease similarity. Additionally we wanted to do so in a manner that was not dependent on specific keywords or ontologies. We demonstrated that LDA seems to be able to represent a disease reasonably and recover its ground truth relations well despite its unsupervised nature. We also showed that such a topic-driven view of disease may be useful in concept normalization and mapping concepts between ontologies. Additionally, since the model lends itself to human interpretation, it has the potential to drive new hypotheses.

## Availability of Supporting Data

This paper focused on the use of both OMIM (http://omim.org) and DO (http://disease-ontology.org). OMIM is the copyrighted material of the Johns Hopkins University, but may be used “ for your personal use, for educational or scholarly use, or for research purposes” and may be downloaded with a proper key here: http://omim.org/downloads. DO is a member of The Open Biological and Biomedical Ontologies (OBO) and may be downloaded freely from their website: http://www.obofoundry.org/cgi-bin/detail.cgi?id=diseaseontology.

Further, the code used in this research is freely available for download from the National Center for Advancing Translational Sciences: https://spotlite.nih.gov/ncats/omimlda.git

**List of Abbreviations Used**
LDA: Latent Dirichlet Allocation
OMIM: Online Mendelian Inheritance in Man
DO: Disease Ontology
NLTK: Natural Language Tool Kit
MeSH: Medical Subject Headings
ROC: Receiver Operating Characteristic
AUC: Area Under the Curve
NER: Named Entity Recognition
PCA: Principal Component Analysis
TF: Term Frequency

## Author’s Contributions

Conceived of and implemented the methodology: JF. Prepared and stitched data: TP and JF. Provided feedback on methodology: RG and NS. Wrote the paper: JF. Edited paper: RG and NS.

## Acknowledgments

We would like to thank the NIH Office of Intramural Training and Education for the Intramural Research Training Award that made this research possible.

